# Spexin and a novel cichlid-specific spexin paralog both inhibit FSH and LH through a specific galanin receptor (Galr2b) in tilapia

**DOI:** 10.1101/853622

**Authors:** Yaron Cohen, Krist Hausken, Yoav Bonfil, Michael Gutnick, Berta Levavi-Sivan

## Abstract

Spexin (SPX) is a 14 amino acid peptide hormone that has pleiotropic functions across vertebrates, one of which is involvement in the brain-pituitary-gonad axis of fish. SPX(1) has been identified in each class of vertebrates, and a second SPX (named SPX2) has been found in some non-mammalian species. We have cloned two spexin paralogs, designated as Spx1a and Spx1b, from Nile tilapia (*Oreochromis niloticus*) that have varying tissue distribution patterns. Spx1b is a novel peptide only identified in cichlid fish, and is more closely related to Spx1 than Spx2 homologs as supported by phylogenetic, synteny, and functional analyses. Kisspeptin, Spx, and galanin (Gal) peptides and their corresponding kiss receptors and Gal receptors (Galrs), respectively, are evolutionarily related. Cloning of six tilapia Galrs (Galr1a, Galr1b, Galr2a, Galr2b, Galr type 1, and Galr type 2) and subsequent *in vitro* second-messenger reporter assays for Gα_s_, Gα_q_, and Gα_i_ suggests that Gal and Spx activate Galr1a/Galr2a and Galr2b, respectively. A decrease in plasma follicle stimulating hormone and luteinizing hormone concentrations was observed with injections of Spx1a or Spx1b *in vivo*. Additionally, application of Spx1a to pituitary slices decreased the firing rate of LH cells, suggesting direct inhibition at the pituitary level. These data collectively suggest an inhibitory mechanism of action against the secretion of gonadotropins for a traditional and a novel spexin paralog in cichlid species.

## Introduction

Spexin (SPX1; also termed neuropeptide Q) was identified first by computational methods [1], and then also by chemical methods [2] in humans. These computational methods have been attempted mostly on the basis of the characteristics of the prohormones from which active neuropeptides are processed. The mature peptide sequence contains 14 amino acids that are flanked by monobasic and dibasic proteolytic cleavage sites. The mature spexin peptide was found to be identical in all tetrapods and elephant shark, and differs in only one amino acid (A^13^T) in piscine species. A paralog of spexin, termed Spx2, was later identified in non-mammalian species [3]. In mammals, Spx was found to participate in inducing stomach contraction [1], inhibiting adrenocortical cell proliferation[4], postnatal hypoxia response [5], cardiovascular and renal modulation[6], nociceptive response [7], fatty acid absorption and weight regulation [8], and diabetes [9]. In teleosts, functional studies of Spx1 mainly focused on its inhibitory role in the regulation of reproduction [10] and food intake [2] [11]. However, a recent study reported that *spx1* knock-out zebrafish exhibited normal reproductive capability but higher food intake than wild type fish, an effect mediated via increased expression of the appetite stimulant, agouti-related peptide AgRP1 [12].

The galanergic neurotransmission system is one of the newest described signaling systems. Today, the galanin family consists of galanin (Gal), galanin-like peptide (GalP), galanin-message associated peptide (GMAP), and alarin, and this family has been shown to be involved in a wide variety of biological and pathological functions [13]. Three different types of galanin receptors have been described so far in mammals: galanin receptor 1, 2, and 3 (GALR1, GALR2, and GALR3) [14]. All of them are members of the G protein-coupled receptor (GPCR) family and act through stimulation of various second messenger systems. The biological activity of GALR1 and GALR3 stimulation is linked to the activity of adenylate cyclase (AC) and cyclic AMP (cAMP) production, and stimulation of GALR2 receptor results in phospholipase C (PLC) activity [14]. Recently, it was reported that SPX is a functional agonist for GALR2 and GALR3 in humans as well as Galr2a and Galr2b in zebrafish [15].

Reproduction is regulated by the hypothalamic-pituitary-gonadal (HPG) axis in all vertebrates. Hypothalamic axons secrete neuropeptides into the gonadotroph cells of the pituitary in order to mediate the expression and secretion of the gonadotropins follicle-stimulating hormone (FSH) and luteinizing hormone (LH) [16]. The first identified neuropeptide that regulates this function was gonadotropin-releasing hormone (GnRH). The hypophysiotrophic type of GnRH regulates gonadotropins by neuroglandular and neurovascular anatomical connections in zebrafish[17; 18]. Recently, a plethora of neuropeptides involved in the control of reproduction were investigated. Most of them are stimulatory neuropeptides, such as neurokinin B [19], neuropeptide Y [20], secretoneurin [21], galanin [22], agouti-related peptide [23], and melanocortin [24]. However, the study of neuropeptides that relay their signal through inhibitory pathways is more challenging, and hence are less studied. The most known inhibitory pathway in fish reproduction is the dopaminergic system [25]. In fish, as in mammals, dopamine D2 receptors transduce their signal through an inhibitory (Gα_i_) signaling pathway [26]. Gα_i_ signaling is involved in a variety of physiologic processes, including chemotaxis, neurotransmission, proliferation, hormone secretion, and analgesia[27].

Nile tilapia is one of the top principal aquaculture species, and is a suitable experimental model fish for reproductive endocrinological research on Perciformes, which is the most recently evolved and largest group of teleost fish that includes many other target aquaculture species. Our objective was to identify spexin in tilapia and clarify its role as a regulator in the HPG axis. This was accomplished by cloning two spexin and six Galr sequences, and performing *in vitro* and *in vivo* studies.

## Materials and Methods

### Fish husbandry and transgenic lines

Sexually mature Nile tilapia (*Oreochromis niloticus*, Lake Manzala strain) were kept and bred in the fish facility unit at the Hebrew University in 500-L tanks at 26°C and with a 14/10 hour light/dark photoperiod regime. Fish were fed daily ad libitum with commercial fish pellets (Raanan fish feed, Israel).

We previously created transgenic tilapia lines by the adoption of a tol2 transposon-mediated approach and Gateway cloning technology [28]. In the current study we used transgenic tilapia in which red fluorescent protein (RFP) expression is driven by the tilapia LHβ promoter, thus labeling LH gonadotrophs. The tagRFP-CAAX cassette, used in the current study directs the fluorescent protein to the cell membranes. The use of tagRFP eliminates the aggregation problems associated with mCherry and results in a more uniform labeling of the cells.

All experimental procedures were in compliance with the Animal Care and Use guidelines of the Hebrew University and were approved by the local Administrative Panel on Laboratory Animal Care.

### Cloning and tissue distribution of tilapia spexin ligands and receptors

The putative spexin 1a and 1b gene sequences were initially identified using tblastn against the tilapia genome (O_niloticus_wgs_v1, Genbank 354508) with zebrafish spexin (XM_005164774) as input, and custom full coding sequence primers were designed for PCR. Primers were designed against the nucleic acid sequences of the six galanin receptors (Galr1a, Galr1b, Galr2a Galr2b, Galr type 1 and Galr type 2) obtained from the Ensembl Genome Browser (http://www.ensembl.org) and the GenBank database (http://www.ncbi.nlm.nih.gov/genbank). Putative transmembrane domains, glycosylation sites and phosphorylation sites were identified using TMHMM v.2.0, NetNGlyc 1.0 [29], and NetPhos 2.0 [30].

Total RNA was extracted from sexually mature female tilapia brain using TRIzol reagent (Life Technologies), and 5 μg was used as template for cDNA synthesis using Smart MMLV reverse transcriptase (Clontech). All cloning PCRs were performed with an initial denaturation at 94 °C for 2.5 min, followed by 30 cycles of denaturation at 94 °C for 30s, annealing at each of the primers’ specific Tm (Table 1) for 30s, and extension at 72 °C for 90s, and a final extension at 72 °C for 10 min using Advantage2 polymerase mix (Clontech); specific primers were designed for cloning the putative spexin ligands and receptors (Table 1). The PCR products were ligated into pCRII-TOPO vector and cloned into competent DH5α *E. coli* cells. Plasmid DNA was isolated from overnight cultures by miniprep columns (QIAgen) and sequenced with T7 and SP6 primers.

**Table 1.**
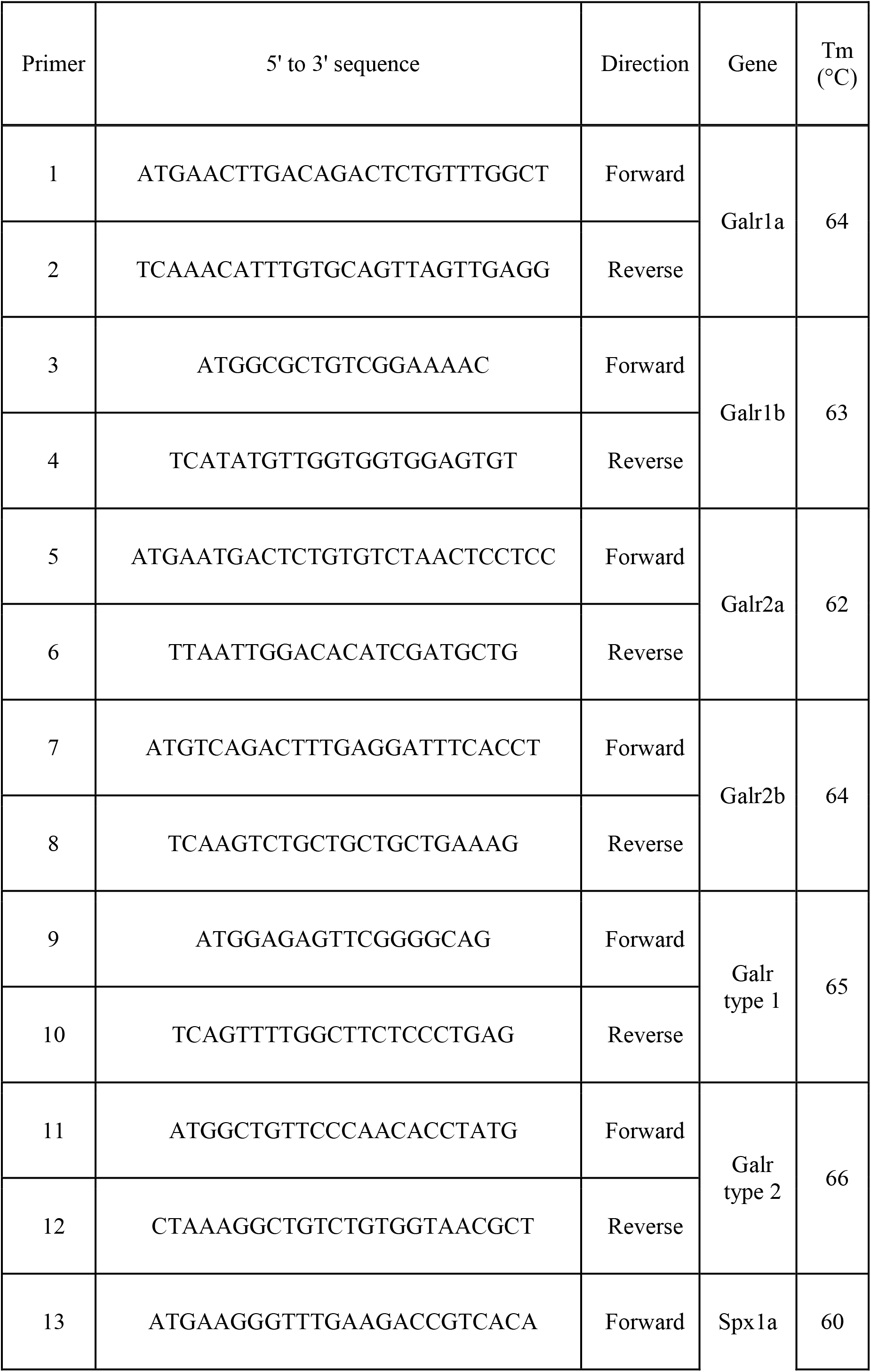
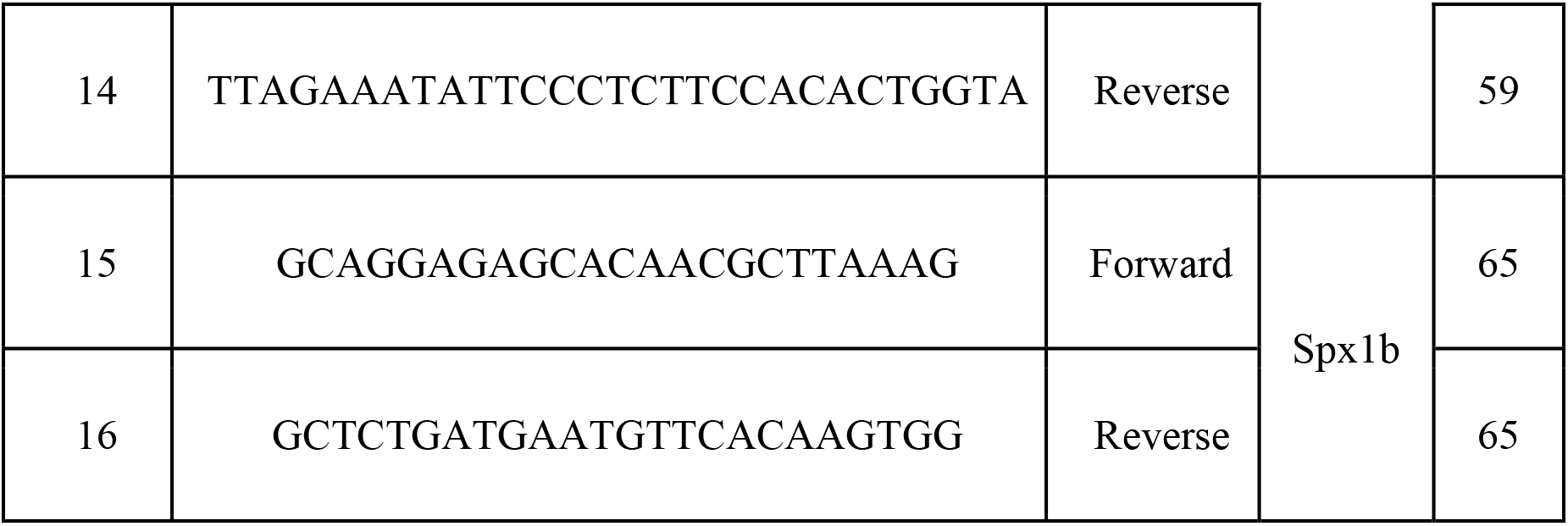
Gene-specific primers used for the identification of tilapia Spx peptides and Galrs. The primers sequences and their melting temperatures used for the PCR are presented.

Cloned tilapia spexin 1a and 1b sequences were submitted to GenBank under accession numbers MN399812 and MN399813 respectively. Cloned tilapia galanin receptors sequences were submitted to GenBank under accession numbers MN326828, MN326829, MN326830, MN326831, MN614146, and MN614147 for Galr1a, Galr1b, Galr2a, Galr2b, Galr type 1, and Galr type 2, respectively.

Tissue samples were collected from three mature male and female tilapia. Total RNA was extracted from each of the following tissues: brain, pituitary gland, spleen, gills, kidneys, muscles, fat, ovaries/testes, retina, heart, caudal and front intestines, and liver. We dissected the brain into three parts, of which the anterior part contains the olfactory bulbs and preoptic area, the midbrain contains the optic tectum and hypothalamus, and the hindbrain contains the medulla oblongata and the cerebellum. cDNA samples were prepared from 2μg of total RNA according to [31]. The tissue expression patterns of Spx1a, Spx1b, Galr1a, Galr1b, Galr2a, and Galr2b in various tissues were analyzed by qPCR with the primer sets described in Table 2. The cycling parameters consisted of preincubation at 95 °C for 10 min followed by 45 cycles of denaturation at 95°C for 10 s, annealing at 60°C for 30 s, and extension at 72°C for 10s, followed by a melting curve analysis (95°C for 60s, 65°C for 60s, 97°C for 1s).

**Table 2.**
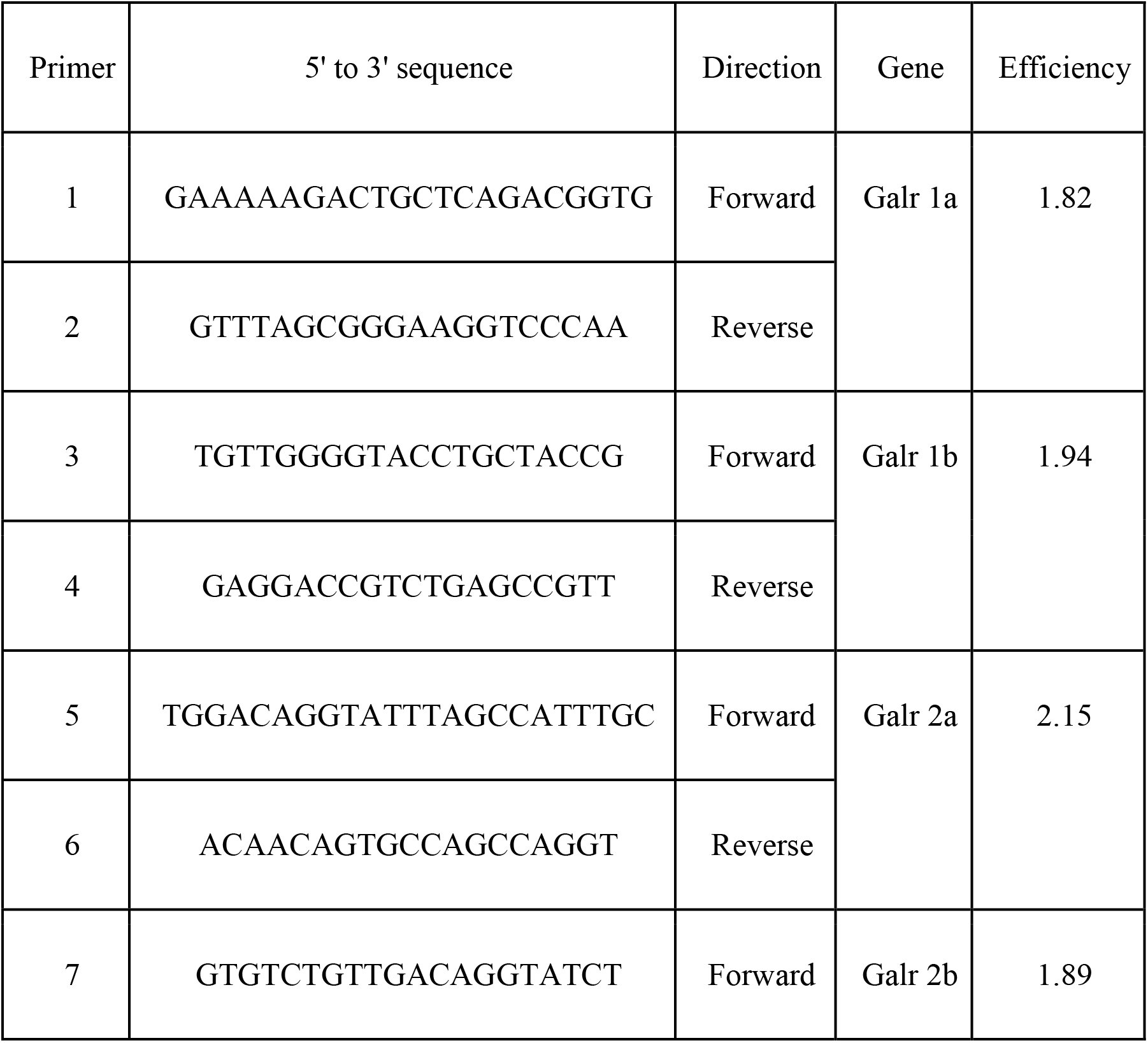
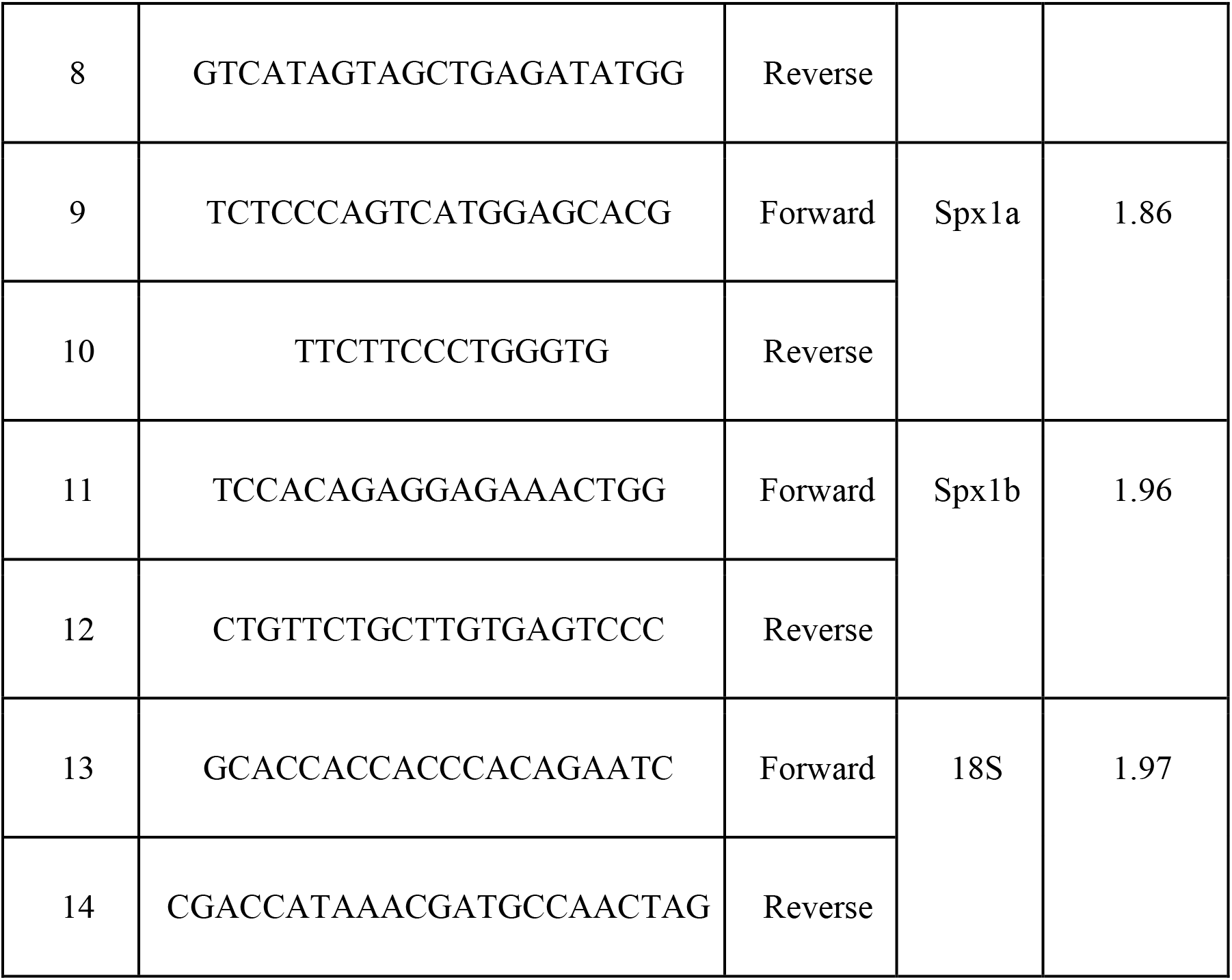
Gene-specific primers used for real-time PCR. The primer sequences and PCR efficiency (10^-1/slope^) are shown.

### Phylogenetic and synteny analyses

All sequences used for spexin and galanin receptors were identified from NCBI and Ensembl databases (Supplementary Table 1) and aligned by MUSCLE using Mega7. Evolutionary analyses were conducted in MEGA7 [32]. The evolutionary history of mature spexin peptide homologs was inferred using the Maximum Likelihood method based on the JTT matrix-based model, and an additional frequency distribution (JTT+F) was utilized for the receptors [33]. The bootstrap consensus trees were inferred from 500 replicates to represent the evolutionary history of the taxa analyzed [34]. The percentage of replicate trees in which the associated taxa clustered together in the bootstrap test are shown next to the branches. Initial tree(s) for the heuristic search were obtained automatically by applying Neighbor-Join and BioNJ algorithms to a matrix of pairwise distances estimated using a JTT model, and then selecting the topology with superior log likelihood value. The analysis involved 34 and 77 amino acid sequences and there were a total of 15 and 264 positions in the final datasets for spexins and Galrs, respectively.

We examined and compared the genomic environment around spexin 1 and spexin2 in zebrafish, Nile tilapia, and Burtoni. Cichlid orthologs were identified by tblasn query using zebrafish genes in NCBI genome browsers, and organized 5’ to 3’ on their respective chromosomes/linkage groups/scaffolds. Accession numbers can be found in Supplementary Table 2.

### Peptide synthesis

Tilapia spexin 1a (NWTPQAMLYLKGTQ-NH2), tilapia spexin 1b (NWTSQAILYLKGAQ-NH2) and human galanin (GWTLNSAGYLLGPHAVGNHRSFSNKNGLTS-NH2) were synthesized by GL Biochem. Peptides were synthesized by the automated solid-phase method by applying Fmoc active-ester chemistry, purified by HPLC to >95% purity and the carboxy terminus of each peptide was amidated. The peptides were dissolved to the desired concentration in fish saline (0.85% NaCl in DDW) for *in vivo* experiments.

### *In vivo* effect of Spxla and Spxlb on FSH and LH release

Adult female tilapia (body weight (BW)= 82.4±21.7g) were injected intraperitoneally with saline, tilapia spexin 1a, or spexin 1b at 10 μg/kg fish (n=8 fish per group). The fish were bled from the caudal blood vessels into heparinized syringes at 2, 4, 8, and 24 h after injection. Blood was centrifuged (3,200 rpm for 30min at 4°C) to obtain plasma samples, which were stored at −20°C until assayed. This time course is according to standard protocols used previously to test the effect of various hypothalamic neuropeptides on circulating levels of LH and FSH in tilapia [35] [36]. Six independent experiments were carried out for the *in vivo* studies.

### ELISA for the measurement of tilapia FSH and LH

Plasma LH and FSH concentrations were measured by specific competitive ELISAs developed for tilapia [37] using primary antibodies against recombinant tilapia LHβ or FSHβ, respectively, and recombinant tilapia LHβα [38] or FSHβα [37] for the standard curves. The sensitivity was 2.43 and 1.52 ng/mL for LH and FSH, respectively. The inter-assay covariance was 14.8% and 12.5%, and the intra-assay variation was 7.2% and 8% for LH and FSH, respectively.

### Determination of brain Spx and Galr expression in fed vs. fasted adult tilapia by qPCR

An *in vivo* experiment where sexually mature, male tilapia (n = 24, BW = 41.7 ± 1. 64) were divided into two groups (one fed ad libitum and the other fasted for 26 days) was previously performed [39]. Total RNA from the brains were taken for real-time PCR expression analyses from the fed and starved groups. The RNA generated from this experiment was used as template for generated cDNA and performing qPCR analysis of brain Spx1a, Spx1b, Galr2a, and Galr2b in this study.

The qPCR cycling parameters consisted of preincubation at 95 °C for 10 min followed by 45 cycles of denaturation at 95°C for 10s, annealing at 60°C for 30s, and extension at 72°C for 10s, followed by a melting curve analysis (95°C for 60s, 65°C for 60s, 97°C for 1s). Reaction conditions were according to [40], and amplification was carried out in Lightcycler 96^®^ (Roche Diagnostic International). Serial dilutions were prepared from a cDNA pool, and the efficiencies of the specific gene amplifications were compared by plotting Ct vs log (template concentration). A dissociation-curve analysis was run after each real-time experiment to confirm the presence of a single product. To control for false-positives, a no reverse transcriptase negative control was run for each template and primer pair. Data were interpreted by the comparative cycle threshold method using 18S as a reference gene [40].

### Second messenger reporter assays

Transient transfection, cell procedures, and stimulation protocols were generally performed as described previously [26] [57]. Briefly, the entire coding sequence of tilapia Galr1a, Galr1b, Galr2a, Galr2b, Galr type 1, and Galr type 2 were inserted into pcDNA3.1 (Invitrogen). Co-transfection of the receptors (3 μg/plate for each Galr) and a cAMP-response element-luciferase (CRE-Luc), serum response element (SRE-Luc; Invitrogen), or Gqi5 reporter plasmid (3 μg/plate) was carried out with TransIT-X2^®^ System (Mirus). The cells were serum-starved for 16 h, stimulated with various stimulants for 6 h, and then harvested and analyzed according to [26] [57]. Experiments were repeated a minimum of three times from independent transfections and each treatment was performed in triplicate wells. EC_50_ values were calculated from dose response curves by means of computerized non-linear curve fitting on baseline-corrected (control) values using Prism version 6 software (GraphPad).

To confirm which of the novel receptors that were cloned in this study transduce effects through the inhibitory Gi signaling pathway, we used a plasmid that contains a chimeric G-protein, Gqi5, in which the five C-terminal amino acids of Gq were changed to those of Gi1,2, (obtained from Addgene, Inc. Cambridge, MA, USA) and has been described previously [41]. Gqi5 is activated by Gi-linked GPCRs, but couples to the effector protein normally activated by Gq, phospholipase C. Since phospholipase C produces IP_3_ and actuates Ca^2+^ release, the inhibitory nature of Gi-linked receptors can be observed as stimulation via the SRE-luc reporter [42].

### Preparation of pituitary slices and electrophysiology

Pituitary slices were prepared according to [43] [44]. Tilapia of both sexes were anesthetized with 2-phenoxyethanol (1 ml/ liter; Sigma-Aldrich Corp., St. Louis, MO) and decapitated. Pituitaries were removed and kept at room temperature in gassed (5% CO_2_) Ringer’s saline containing (in mM): 124 NaCl, 3 KCl, 2 CaCl_2_, 2 MgSO_4_, 1.25 NH_2_PO_4_, 26 NaHCO_3_, and 10 glucose (pH 7.35) up to 5 min before being transferred to 2% agarose low gelling temperature (Sigma) at 32°C. The agarose with the pituitary was placed in the −20°C for a short time until the agarose solidified. The block was glued to the stage of a vibratome (series 1000, Vibratome Co., St. Louis, MO), and 150 μM slices were transferred to cooled gassed Ringer’s saline.

For electrophysiological recordings, slices were gently transferred to a chamber attached to the stage of an upright microscope (Axioskop FS, Zeiss, Oberkochen, Germany) continuously superfused with Ringer’s saline at room temperature. Endocrine cells were viewed with a 40x, 0.8 numerical aperture, water immersion objective lens (Olympus, Munich, Germany). Patch pipettes were pulled from borosilicate glass capillaries (Hilgenberg, Maisfield, Germany) on a Narishige PP83 puller. Membrane currents were recorded using an on-cell patch technique - cell-attached recording in which a patch electrode is attached to the cell, but the membrane is not broken. The standard pipette solution contained (in mM): 135 potassium gluconate, 2 MgCl_2_, 1 CaCl_2_, 11 EGTA, 3 ATP (magnesium salt), and 10 HEPES (potassium salt), pH 7.25. Only fluorescently labeled cells from the pituitary were recorded. An Axoclamp-2B amplifier (Axon Instruments, Union City, CA) was used in Bridge mode. Experiments were controlled by USB-6341 data acquisition board (National Instruments, USA) and WinWCP V5.3.7 (Strathclyde Electrophysiology Software, UK). Data were analyzed using PClamp 10 (Axon Instruments) and Microcal Origin 6.0 software.

### Statistical analyses

Data are presented as means ± SEM. All samples had equal variance, as determined by an unequal variance test performed using JMP 7.0 software. The significance of differences between group means of plasma gonadotropin levels were determined by ANOVA, followed by Tukey’s test using Graph-Pad Prism 5.01 software (San Diego, CA). EC_50_ values of the receptors assays were calculated using log treatment vs. luciferase intensity on a nonlinear regression curve using Prism.

## Results

### Cloning and tissue distribution of spexin1a, 1b, and gal receptors in tilapia

The cloned full coding sequence of tilapia spexin 1a and 1b were 363 and 315 base pairs and coded preprohormones of 120 and 104 amino acids, respectively. Both mature tilapia spexin peptides were presumed to be 14 aa based on conserved peptide-flanking monobasic and dibasic cleavage sites (Figure 1). The amino acid sequence of Spx1b differs from that of orthologous piscine Spx1a at positions 4 (Pro to Ser), 7 (Met to Ile), and 13 (Thr to Ala). Both Spx1a and Spx1b are likely amidated at their C-termini due to the GRR motif. The amino acid sequence of Spx2 differs from that of Spx1 at positions 3 (Gly vs Thr), 6 (Ser vs Ala), 13 (Arg vs Thr (piscine species) or Ala (shark and tetrapods)), and 14 (Tyr or His vs Gln) (Figure 1).

**Figure 1.**
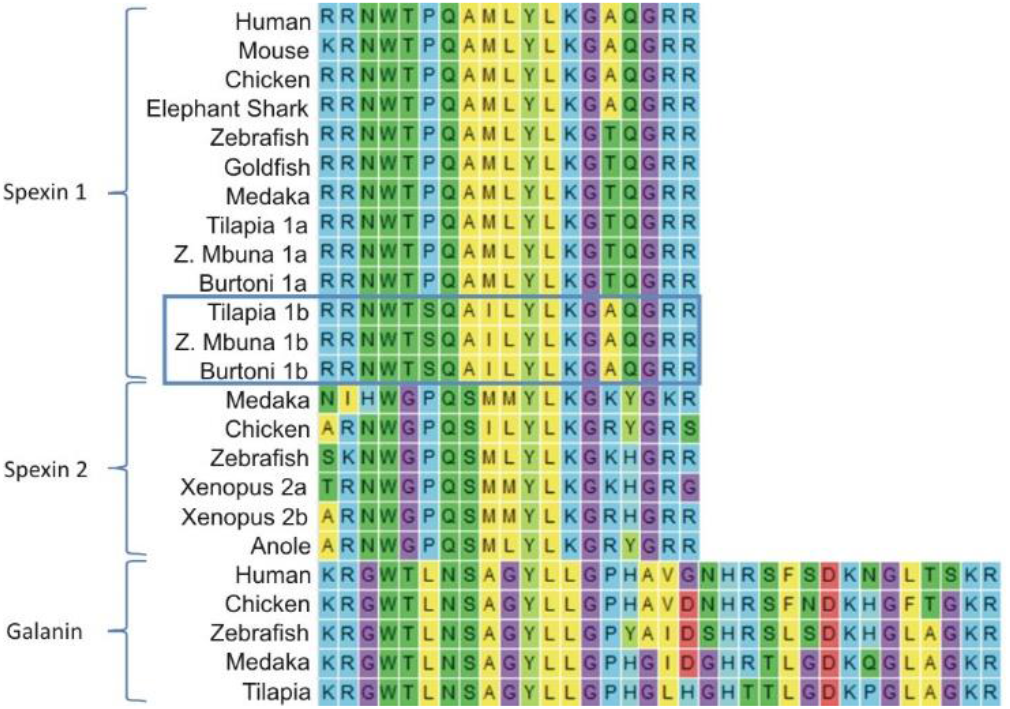
Multiple sequence alignment of mature vertebrate spexin1, spexin2, and galanin peptides with flanking monobasic or dibasic cleavage motifs. The last three amino acids of spexin sequences (GRR) were aligned to the end of the galanin sequences, but manually moved for brevity. Cichlid spexin 1a and 1b sequences are labeled to differentiate them, and spexin1b are boxed.

Six different Galrs were found in the tilapia genome and their corresponding coding regions were cloned and sequenced. According to their positions in the phylogenetic tree, they were named Galr1a, Galr1b, Galr2a Galr2b, Galr type 1 and Galr type 2. They encoded proteins of 350, 357, 325, 379, 344 and 379 amino acids for Galr1a, Galr1b, Galr2a Galr2b, Galr type 1, and Galr type 2, respectively. All seven-transmembrane receptors contain 1-2 potential sites for N-linked glycosylation (NXT/S) in the N-terminal region, and 1-3 serine residues for potential protein kinase C phosphorylation (SSR/SKK/SKR) (Supplementary Figure 3).

To shed light on the potential physiological roles of spexins/Galrs in tilapia, we examined their mRNA tissue distribution by real-time PCR (Supplementary Figure 1). The two spexins and their related receptors exhibit overlapping and distinctive patterns of expression in various tissues. Spx 1a expression was detected primarily in the midbrain and ovary, whereas Spx 1b was detected in pituitary, kidney, and all brain parts, but mostly in the anterior brain. Spx 1b expression levels were a few orders of magnitude higher than that of Spx1a relative to 18S ribosomal subunit expression. The expression patterns of Galr 1a and Galr2a were similar, where both were detected in most of the tissues with the highest expression observed in the kidney and head kidney. However, Galr1b was detected mostly in the anterior gut and anterior brain, with lower expression found in all other tissues except the gills. High expression of Galr2b was detected in the anterior and midbrain relative to the other tissues, where virtually no expression was detected.

### Evolutionary history of spexins and gal receptors

The phylogenetic analysis showed that the vertebrate SPX sequences fall into two distinct clades, with both cichlid spexins grouped with Spx1 (Figure 2A) and all Spx2 orthologs grouped together in a distinct clade. However, the second tilapia spexin paralog was more phylogenetically related with SPX1 than with SPX2 homologs. We also found that other cichlids, like zebra mbuna and Burtoni, contain two spexin paralogs that are more similar to orthologous spexin 1 sequences. Thus, we named the two cichlid spexins Spx1a and Spx1b.

**Figure 2.**
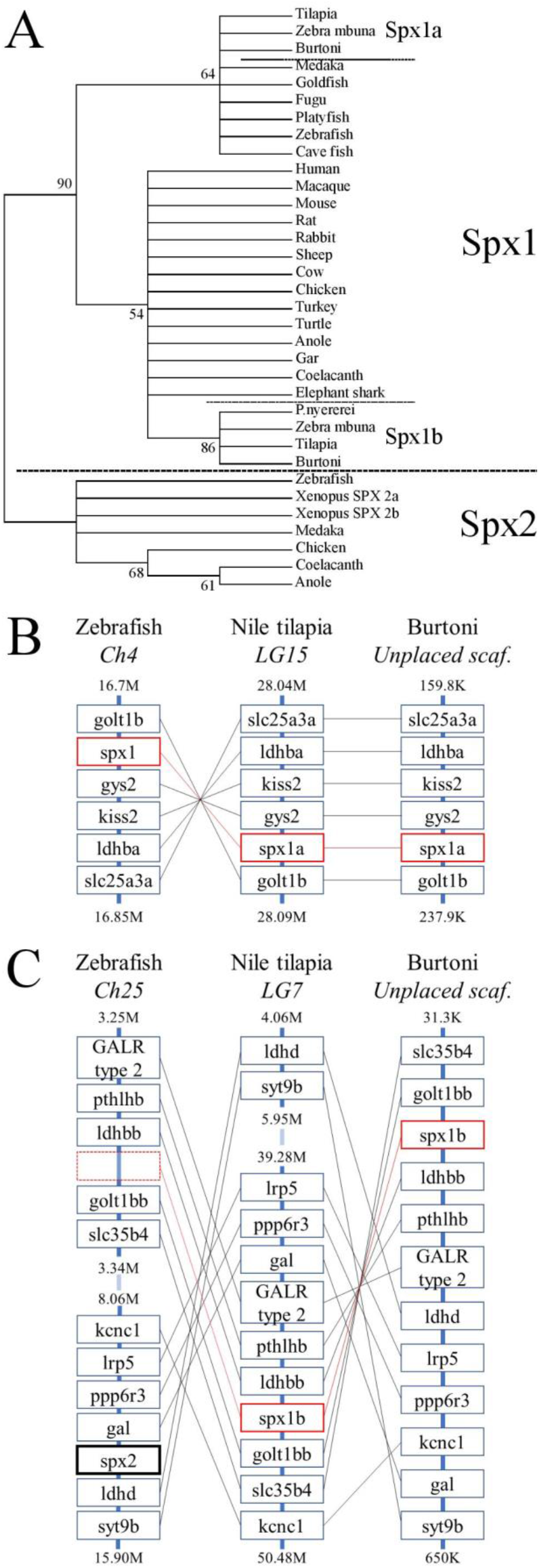
Evolutionary history of spexin sequences. **A.** The phylogenetic tree of mature vertebrate spexin paralogs was generated by the maximum likelihood method and the percentage of replicate trees in which the associated taxa clustered together in the bootstrap test (500 replicates) are shown next to the branches; bootstrap scores lower than 50% are condensed. **B.** The genomic environment of *spx1* and **C.** *spx2* in zebrafish, Nile tilapia, and Burtoni. Spx1a and spx1b are surrounded in red boxes (broken to emphasize lack of a spx-related gene in zebrafish) and Spx2 has a bolded black box. Chromosomes (Ch)/ linkage groups (LG)/ scaffolds (scaf.) are organized 5’-3’ and genomic location (in million (M) or thousand (K) base pairs) is indicated above and below each column. Note that the range of zebrafish Ch25 and tilapia LG7 are broken for brevity.

The genomic organization of spexin paralogs between zebrafish, Nile tilapia, and Burtoni was determined. The genomic environment for *spxla* was identified on zebrafish chromosome 4, Nile tilapia LG15, and Burtoni on an unplaced scaffold, and was highly conserved (Figure 2B). *Spxlb* was identified on Nile tilapia LG7 and an unplaced scaffold in Burtoni between *golt1bb* and *ldhbb. Spx2* was identified on zebrafish chromosome 25 between *gal* and *ldhd* (*Figure 2C*).

To determine the phylogenetic identity of the cloned Gal receptors we performed a phylogenetic analysis consisting of vertebrate Galr and Kissr sequences (Figure 3). The Galr and Kissr sequences formed two distinct clades. The Galr clade was subdivided into Galr1 and Galr2/3 clades, which, respectively, further formed distinct groups of Galr1a and Galr1b, and Galr3, Galr2a, and Galr2b. Tilapia Galr type 1 and 2 and medaka Galr type 1 belong to a sister group of Galr1.

**Figure 3.**
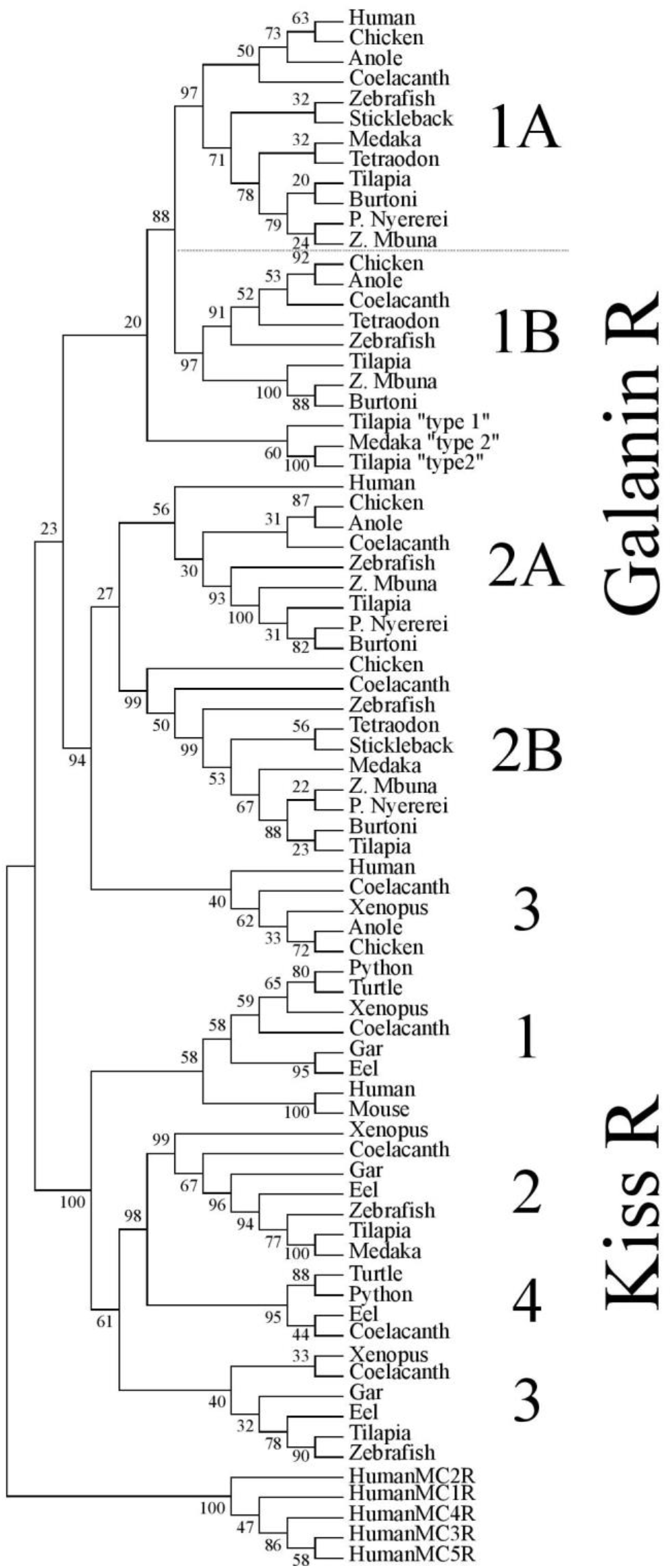
Phylogenetic analysis of vertebrate galanin and kisspeptin receptors. The phylogenetic tree was generated by the maximum likelihood method and the percentage of replicate trees in which the associated taxa clustered together in the bootstrap test (500 replicates) are shown next to the branches. This cladogram was rooted with human melanocortin receptors (MCR). A solid line subdivides the Galr and KissR clades, and subsequent subdivisions within these clades are separated by dashed lines.

### Tilapia spexins suppress FSH and LH release *in vivo*

We next aimed to evaluate the biological effects of the two spexin paralogs on the release of tilapia gonadotropins. After a single intraperitoneal administration of 10 μg/kg fish Spx1a, both FSH and LH plasma levels significantly decreased after 120 min, compared with the levels observed at 0 min, and stayed at this lower level even after 24 h (Figure 4A, C). Injection of Spx1b caused a gradual decrease in plasma levels of FSH and LH, with lower levels shown after 4 h (Figure 4B, D). In the control groups, plasma gonadotropin levels did not change for the whole experimental period.

**Figure 4.**
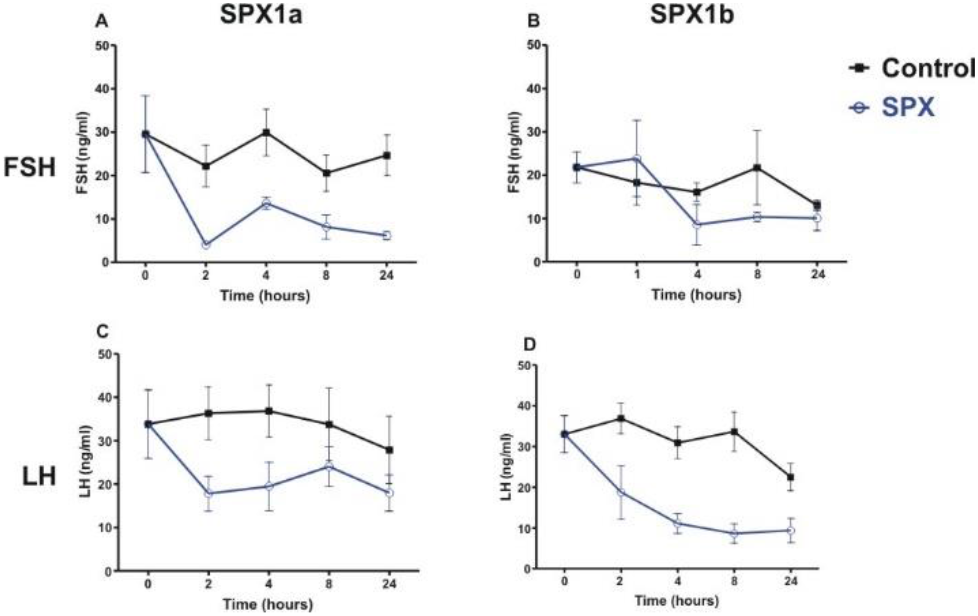
Spexins inhibit LH and FSH *in vivo*. Adult female tilapia were injected with Spx1a, Spx1b, or saline and blood was sampled up to 24 hours. Plasma FSH (**A,B**) and LH (**C,D**) were determined by ELISA. Control values are shown in black, and spexin treatment values are shown in blue. N=8 fish per group. Data are represented as mean FSH or LH (ng/ml) ± SEM.

### Fasting lowers Spexin and Galr2 mRNA expression in adult tilapia

Most of the documented knowledge relates spexin to metabolic processes or feeding behaviors. Even though our study primarily focused on reproduction, we aimed to determine differences in spexin expression in adult tilapia that were fasted or fed for 26 days. Spx1a and Spx1b mRNA expression was significantly lower in the fasted vs. fed fish. Galr2a expression was significantly lower in fasted fish than in fed fish, and Galr2b expression not affected (Figure 5).

**Figure 5.**
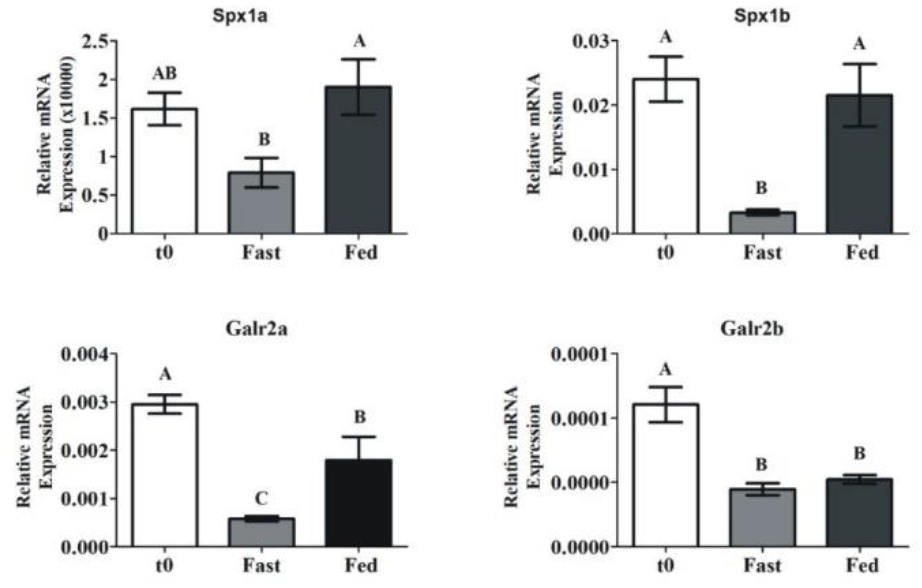
Brain spexin and Galr mRNA expression in response to fasting. Sexually mature tilapia were fed *ad libitum* (Fed), or fasted (Fast) for 26 days. Results were normalized to the amount of *18S* by the comparative cycle threshold method. N = 24. Values are given as mean relative expression ± SEM. Different letters indicate statistically significant differences between groups (*p* < 0.05).

### Activation and signaling of tilapia galanin receptors

All six cloned, full-length tilapia Galrs were sub-cloned into the pcDNA3.1 expression vector and co-transfected into COS7 cells with a luciferase plasmid containing either cAMP response element (CRE), serum responsive element (SRE), or Gqi5 to evaluate their signal transduction pathway(s) upon treatment with tilapia Spx1a, Spx1b, or human galanin (Figure 6). Since each Galr subtype is coupled to different G proteins (Gα_i_ for GalR1 and GalR3; Gα_q/11_ for GalR2) [14], we tested all the receptors using three different signal transduction pathways: CRE-luc for the activation of PKA/cAMP, SRE-luc for the activation of PKC/Ca^2+^, and Gqi5/SRE-luc for further verification of inhibition through Gα_i_. The 5 aa at the carboxyl-terminus of Gqi5 are sufficient to enable interaction with Gα_i_-coupled receptors [41]. Ligand stimulation of a Gα_i_-coupled receptor is expected to trigger the following cascade: Gqi5, phospholipase C, protein kinase C, SRE-luciferase transcription, luciferase activity, light emission. For verification of the activation of Gqi5 trough Gα_i_, we used the tilapia dopamine D2 receptor [26][57]. A dose-dependent increase in luciferase activity confirmed that tilapia D2 receptor relayed its activity trough the inhibitory Gi (Supplementary Figure 4).

**Figure 6.**
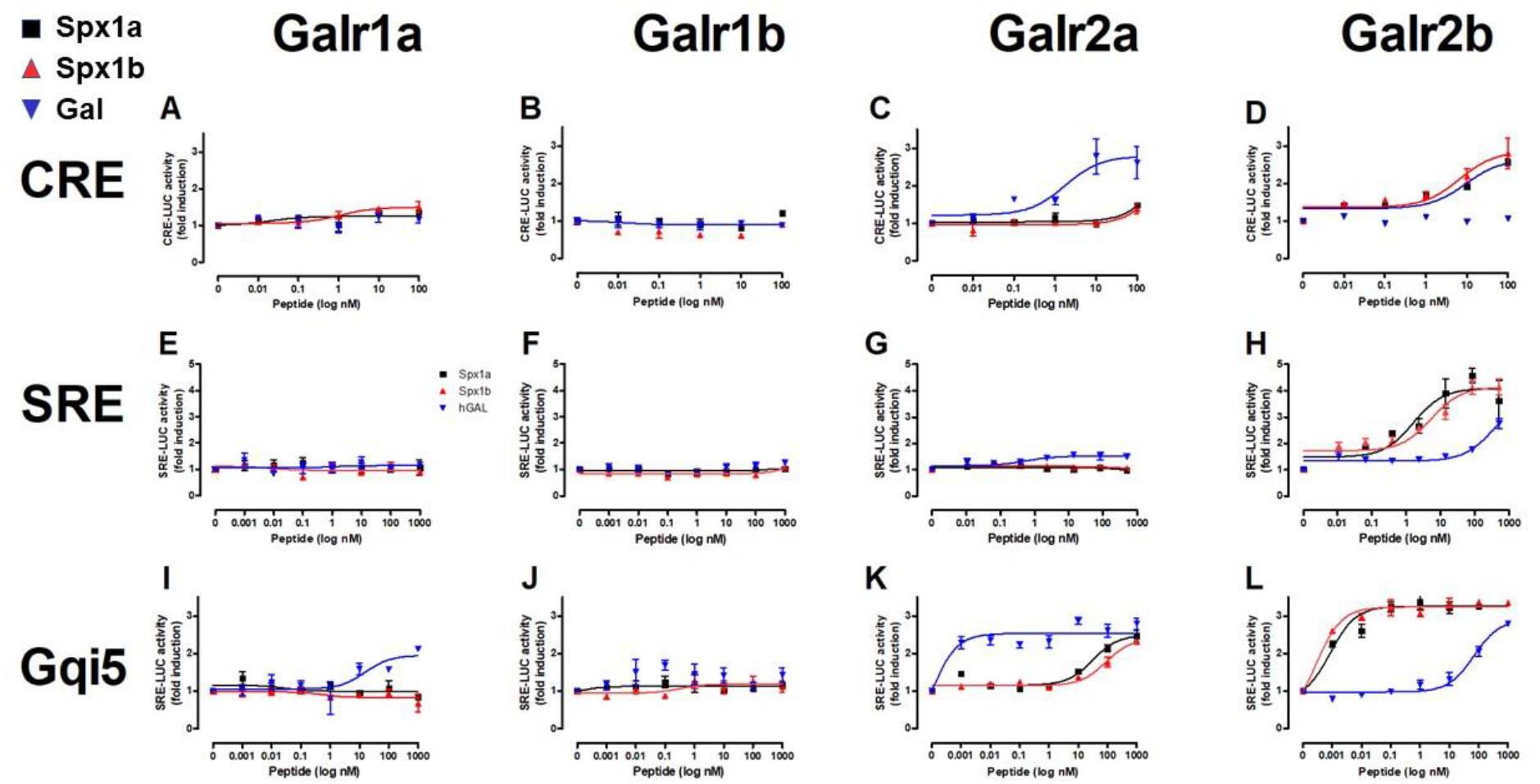
*In vitro* reporter assays for tilapia Galanin receptors. Tilapia Galrs were cotransfected with either CRE-luc (**A-D**), SRE-Luc (**E-H**), or Gqi5 (**I-L**) into Cos7 cells and treated with either Spx1a (black squares), Spx1b (red triangles), or human galanin (Gal; blue upside-down triangles). Data are represented as mean ± SEM fold-change in luciferase activity over basal (no treatment).

Tilapia Spxs or hGAL did not increase CRE-luc or SRE-luc levels in cells expressing Galr1a or Galr1b, but hGAL activated Gqi5 signaling via Galr1a. Increasing concentrations of Gal resulted in significant increases in CRE-luc, SRE-luc, and Gqi5 activity in cells expressing Galr2a. Stimulation by high concentrations of both tilapia Spx paralogs resulted in an increase in Gqi5 in cells expressing Galr2a. Exposure to increasing concentrations of either Spx resulted in a dose-dependent increase in SRE-luc, CRE-luc, and Gqi5 activity in cells expressing Galr2b. Both Spx1a and Spx1b were also very efficient in transcription of SRE-luc in cells expressing Gqi5, suggesting that Galr2b relays its signal through Gi (EC_50_=1nM). The EC_50_ values of Spxs and Gal for each receptor are summarized in Supplementary Table 3.

### Effect of spexin1a on the electrophysiological activity of pituitary LH cells

Following the results that spexins activate Galr2a and 2b, of which the former was also found to be expressed in the pituitary of tilapia, we aimed to examine whether spexin affects LH gonadotroph electrical activity. To address this question, we used an electrophysiological method on disconnected pituitary slices to keep the cells in their natural organization, but without signals from the brain. We measured the frequency of action potentials before, during, and after application of 100nm Spx1a. Exposure to Spx1a decreased the spike rate, which slowly increased back to basal frequencies when spexin was washed out from the recording chamber (Figure 7).

**Figure 7.**
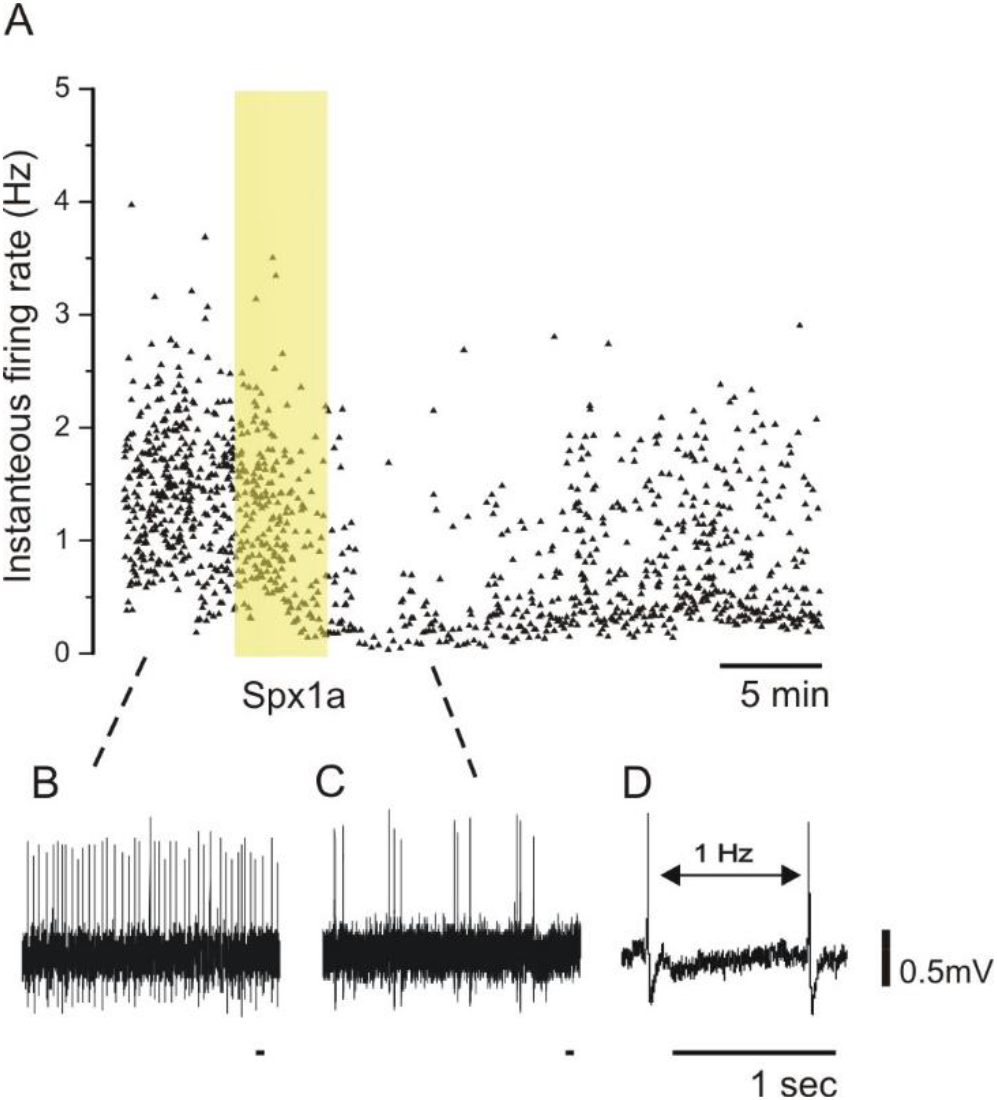
Effect of spexin1a on firing rate of LH cells in mature tilapia. Transgenic tilapia (LH-RFP) pituitaries were dissected for electrophysiological recording, and a cell-attached configuration was used to monitor action potentials without breaking the cell membrane. The pituitary slice was briefly exposed to Spx 1a during the recording (blue bar). **A**. A graphic representation of firing-rate before (**B**), during (**C**), and after spexin application. Each dot represents a action potential along the time scale (X-axis) and the instantaneous frequency from the action potential before (Y-axis). Note that the firing-rate decreased after Spx1a was applied, and after a few minutes the spike rate slowly rises back to the control level. **D**. Two spikes with an example of the firing-rate measurement.

## Discussion

Spexin is a highly conserved peptide hormone that may have pleiotropic functions in different vertebrate species. Cloning of two spexin paralogs and six galanin receptors permitted phylogenetic, synteny, and functional analyses in Nile tilapia, supporting the involvement of the spexin/galanin receptor system in fish reproduction and metabolism. Evolutionary analyses support that cichlid species possess a novel form of Spx1, named Spx1b, but not Spx2 like in some other vertebrates. Demonstration of Spx1a and Spx1b involvement in reproduction was shown *in vivo* and *in vitro*, and starvation limited the expression of both paralogs. Despite their differential tissue distribution and expression profiles, both Spx1a and Spx1b had similar biological effects, which are likely mediated through galanin receptor 2b.

Phylogenetic and synteny analyses coupled with vertebrate ancestral chromosome (VAC) reconstruction supported that *kiss, gal*, and *spx* genes were located adjacent to each other on VAC D and individually arose by local tandem duplication prior to the first whole genome duplication event [15]. This genomic proximity was conserved after two rounds of whole genome duplication (1R, 2R) and is observed in modern vertebrate genomes, where *spx1* is found near *kiss2*, and *spx2* is found near *gal*. Our synteny analysis supported this organization for Nile tilapia and Burtoni *spx1*, but not *spx2*. We initially had thought that the spexin-like gene flanked by *golt1bb* and *ldhbb* in tilapia was *spx2*, but this did not correspond with the genomic organization of zebrafish *spx2*, which is flanked by *gal* and *ldhd*. The genomic distance between Nile tilapia *gal* and *ldhd* is >34 million base pairs, and *spx1b* lies outside of this syntenic block between *golt1bb* and *ldhbb*. No zebrafish spexin-like gene was found between *golt1bb* and *ldhbb*, suggesting that this tilapia and Burtoni spexin paralog is not *spx2*. Additionally, Spx1b has 85-93% (12-13/14aa) sequence conservation across all vertebrate Spx1 peptides, but only 57-71% (8-10/14aa) conservation with the variable and meager Spx2 sequences. Phylogenetic analysis of mature spexin peptide sequences shows that cichlid Spx1b forms a sister clade to vertebrate Spx1, and is not placed with other teleost Spx1 homologs. Due to the highly conserved nature of short peptide sequences and a 20% divergence in amino acid sequence identity from Spx1, we propose that Spx1b is a novel spexin peptide found only in cichlid species.

Spexin has been implicated in regulating the hypothalamic-pituitary-gonad (HPG) axis in some fish species. Our *in vivo* experiments support that not only Spx1a, but also the novel form, Spx 1b, inhibits LH and FSH release in tilapia. The first report on fish spexin in zebrafish and goldfish identified high and dynamic expression in the brain throughout the reproductive stages [10]. Spexin inhibited LH release in goldfish pituitary cultures and *in vivo*, and an estradiol feedback mechanism on hypothalamic spexin expression was observed. Ovariectomized goldfish had increased hypothalamic spexin expression, and estrogen replacement returned expression to basal levels [10]. A similar effect was seen in the spotted scat, where estradiol decreased spexin expression in a dose-dependent manner *in vitro* [45]. However, zebrafish *spx^−/−^* knockouts displayed normal reproductive phenotypes, so spexin may only play a direct role in regulating the reproductive axis in certain fish species [12]. With that being said, knockout models of reproductive hormones in fish species tend to reproduce normally (reviewed in [46]). There are few reports on the role of spexin in reproduction, all of which have been carried out in fish; additional studies in non-piscine species are required to determine if spexin has a functionally conserved role along the HPG axis.

A distinctive feature of the teleost pituitary gland is that LH and FSH are synthesized by different gonadotrope cells, which suggests a functional and differential regulation of gonadotropin release mechanisms. The cells of the teleost pituitary are distinctly organized and receive hypothalamic signals directly by nerve innervation as well as via neurovasculature [18]. Teleost LH cells form dense networks throughout the pars distalis of the adenohypophysis, and FSH cells form small clusters that are more broadly distributed throughout the pars distalis [17]. LH cells, but not FSH cells, are functionally coupled, as shown by perfusion of pituitary fragments exposed to GnRH with gap-junction blockers and by a patch-clamp technique [17]. These organizational characteristics of the teleost pituitary complicate our understanding about the various ways by which the gonadotropes decode stimulatory and inhibitory inputs. The principal stimulatory and inhibitory factors for LH release are gonadotropin-releasing hormone (GnRH) and dopamine (DA), respectively, but less is known about FSH release. LH and FSH cells are regulated by more than 20 neurohormones, but only a small portion of them function as inhibitory factors [46]. In fish, the dopamine receptor (D2-R) is expressed on LH cells [47] and on GnRH neurons [25], suggesting that direct and indirect mechanisms of LH release inhibition exist. Gonadotrophs are excitable cells, and action potential frequency of LH and FSH cells were shown to be influenced by GnRH exposure [48]. We performed electrophysiological measurements on LH cells in pituitary slices and observed that Spx1a decreases the action potential frequency, suggesting that spexin decouples the LH gonadotrope functional network directly.

Spexin mRNA expression levels may also be correlated with reproductive development. Increases in LH and FSH mRNA expression is seasonal and corresponds to gonadal development [49] [50]. In goldfish, hypothalamic Spx1 expression significantly decreased as the breeding season progressed and GSI increased [10], potentially contributing to progressive LH release. A similar decrease was seen in the orange-spotted grouper over the course of oogenesis, where the highest hypothalamic Spx1 expression was observed prior to oocyte primary growth [11], and in the spotted scat, expression significantly decreased over the course of vitellogenesis [45]. In zebrafish, however, Spx1 expression in the brain gradually increased from the ovarian primary growth stage, peaked in early-mid vitellogenesis, then returned to primary growth levels during final maturation [10]. Given that Spx1 is negatively regulated by estrogen [10], which increases in circulation until final oocyte maturation, it seems reasonable that Spx expression correlates with reproductive development. Additionally, Spx1 treatment affects the expression of other reproductive factors, such as increasing gonadotropin inhibitory hormone (GnIH) and GnRH-III expression and decreasing GpHα and FSHβ expression in a sole [51].

Spexin has wide tissue distribution patterns in tetrapods and fish, suggesting that spexin has pleiotropic biological consequences in feeding behavior, metabolism, nociception, cardiovascular function, muscle motility, stress, depression, and anxiety [3]. The localization of spexins at the cellular level inform these functions. In tilapia, the two spexin paralogs exhibited wide differential tissue distribution patterns, where Spx1a was primarily expressed in the midbrain (containing the hypothalamus in our preparations), and Spx1b was primarily expressed in the anterior brain, which contains the preoptic area. Similarly, in a detailed report of spexin neuron circuitry in zebrafish, Spx1 expression was restricted to the midbrain tegmentum and the hindbrain and Spx2 expression was found in the preoptic area, however neither was detected in the hypothalamus [52]. In goldfish, Spx1 was identified in the hypothalamus, thalamus, and medial longitudinal fasciculus, however immunoreactivity was detected using heterologous antibodies that may have recognized both Spx1 and Spx2 [10]. Even though our tissue distribution analysis is crude and unable to implicate any detailed biological function from expression, these data support the conservation of Spx1 expression, which differs from Spx1b localization. In order to determine if the neuronal circuits of spexins are conserved in fish, methodologies for determining specific spexin localization (IHC with custom antibodies, fluorescent *in situ* hybridization, etc.) are needed from additional species.

Spexin plays a role in feeding behavior and metabolism in fish and mammals (reviewed in [3]). In the orange-spotted grouper [11], half-smooth tongue sole [45], and spotted scat [45], starvation increased Spx1 expression. We observed a decrease in brain Spx1a, Spx1b, and Galr2a expression in fasted adult tilapia. Brain Spx1 expression was also decreased in unfed Ya-fish [53]. In goldfish, Spx1 injections decreased surface feeding and increased food rejection, and increased circulating Spx1 and brain and liver Spx1 mRNA expression after feeding [2] [54]. This effect was also observed in *spx1^−/−^* zebrafish, which had higher food intake compared to wildtype fish, and Spx1 administration suppressed agouti-related peptide (AgRP) expression [12]. In support of this mechanism of satiety control, central administration of Spx1 in goldfish caused a decrease in expression of appetite stimulants (NPY, AgRP, and apelin) and increased expression of anorexigenic factors (proopiomelanocortin (POMC), cocaine- and amphetamine-regulated transcript (CART), cholecystokinin (CCK), melanin-concentrating hormone (MCH), and corticotropin-releasing hormone (CRH)) in different parts of the brain [2]. Furthermore, intracerebroventricular injection of Spx1 inhibited feeding behavior induced by NPY and orexin. In addition to appetite factors, insulin was shown to increase circulating levels of Spx1 and its expression in the telencephalon, hypothalamus, and optic tectum [54]. Therefore, local and central Spx1 appears to be an integral part of appetite control and energy homeostasis in fish.

Spexin mediates its effects by activating an inhibitory G-protein (Gα_i_) via galanin receptor (GalR) 2/3. Like the kisspeptin, galanin, and spexin peptides, the kisspeptin and galanin receptors are related. Ancestral forms of KissR, GalR1, and GalR2/3 were identified on different VACs with 1R and 2R expanding each clade, giving rise to four KissR (KissR1, 3, 2, 4), two GalR1 (1a, 1b), two GalR2 (2a, 2b) and GalR3 paralogs [15]. Our phylogenetic analysis supports that of Kim *et al*. (2014), showing that GalRs are divided into two major clades, namely GalR1 and GalR2/3, with KissRs forming a distinctive sister clade. We cloned two additional Galrs based on predicted NCBI sequences named Galr type 1 and type 2, which formed a sister group to GalR1 sequences, but were not activated by galanin or spexin (Supplementary Figure 2). It has been previously shown that the cognate receptor(s) for Spx1/Spx2 are GalR2/3, whereas GalR1 is the cognate receptor for galanin; teleosts do not possess GalR3 [15]. Our second messenger reporter assays revealed that tilapia Spx1a and Spx1b activated Gα_s_ (via CRE), Gα_q_ (via SRE), and Gα_i_ (via Gqi5) signaling pathways through Galr2b, but not Galr1a or Galr1b. Galanin was more efficacious than Spx1a or Spx1b in activating Gα_s_ and Gα_i_ via Galr2a, but did not activate Gα_q_ (Figure 8). Determining inhibitory effects via Gα_i_ is challenging, so we utilized a reporter plasmid that permits Gα_i_-coupled receptors to stimulate PLC [41]. Human, Xenopus, and zebrafish GalRs were activated in a similar manner using an alternative expression system [15]. GalR2 displays differential signaling preferences to either Gal or Spx, which induce different conformational changes in the receptor and bias intracellular signaling cascades [55]. Spx quickly dissociated after G-protein signaling, whereas Gal binding was more stable and promoted arrestin-dependent receptor internalization. It has been shown that galanin and spexin have an inverse relationship in regards to LH release (reviewed in [15]), perhaps due in part to the receptor displaying biased agonism in order to decipher endogenous hormone signaling.

**Figure 8.**
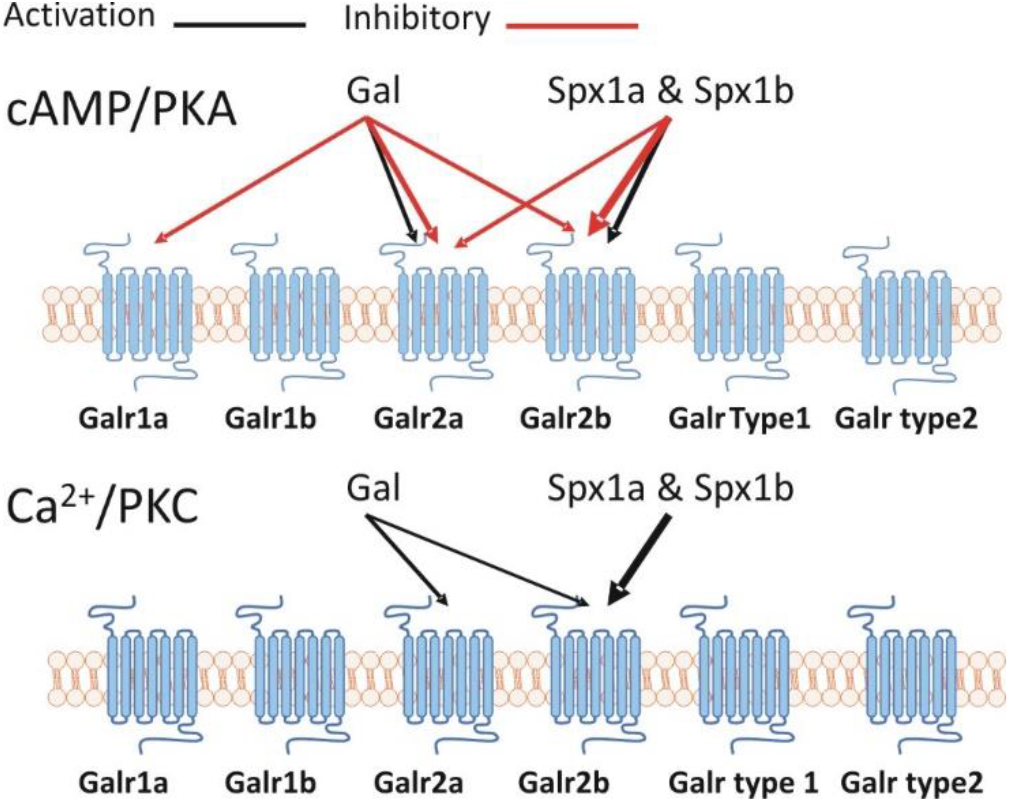
A schematic representation of the Spx/Galr system in tilapia.

We have shown evolutionary and functional evidence that cichlid fish possess two paralogs of Spx1 but not Spx2, and that tilapia Spx1a and Spx1b can inhibit LH and FSH secretion via Galr2b. Why cichlids have evolved a second form of Spx1 and lost Spx2 remains unknown. One in ten teleosts are cichlids, making them the most species-rich group of vertebrates and excellent genetic models of adaptive radiation [56]. Additionally, Nile tilapia is a major globally aquacultured species. Further research into the potential for manipulation of growth and reproductive processes with spexin or galanin receptor agonists/antagonists is warranted.

## Acknowledgements

The research was funded by the Israel Science Foundation (ISF) no. 1540/17. KNH was supported by BARD, the United States - Israel Binational Agricultural Research and Development Fund, Vaadia-BARD Postdoctoral Fellowship Award No. FU-561-2017.

We would like to thank Dr. Michal Shpilman for her technical assistance.

## Author Contributions Statement

YC, KH, and BS prepared and wrote the manuscript. YC, KH, and YB curated and analyzed all data. MG and BS acquired funding, and supervised and conceptualized the manuscript. All authors contributed to manuscript revision, read and approved the submitted version.

## Contribution to the Field Statement

Reproduction is regulated by a hierarchical hormonal signaling cascade via the brain-pituitary-gonadal axis in all vertebrates. Many neurohormones release from the brain have been identified to stimulate the release of pituitary gonadotropins (follicle stimulating hormone and luteinizing hormone), which stimulate reproductive developmental processes (I.E. gametogenesis) in the gonads. A plethora of neuropeptides involved in the control of reproduction have been investigated over the last decades, and most of them are stimulatory neuropeptides. However, the study of neuropeptides that relay their signal through inhibitory pathways is more challenging. Spexin is a highly conserved 14 amino acid peptide hormone that may have pleiotropic functions in different vertebrate species, but was shown to inhibit pituitary gonadotropin release in fish. Our work supports that the conserved spexin peptide, as well as a novel cichlid-specific spexin, directly inhibits gonadotropin release in tilapia. Understanding the stimulatory and inhibitory factors that regulate fish reproduction is critical to aquaculture success, as well as our basic understanding of comparative reproductive endocrinology in vertebrates.

