## Supplementary Table 3 for "Spexin and a novel cichlid-specific spexin paralog both inhibit FSH and LH through a specific galanin receptor (Galr2b) in tilapia"

**Table 3. EC50 values (log nM) of tilapia galanin receptors.**

| **Receptor/**  **Ligand** | **Pathway** | **GALR1a** | **GALR1B** | **GALR2A** | **GALR2B** | **GALR type 1** | **GALR type 2** |
| --- | --- | --- | --- | --- | --- | --- | --- |
| **SPX1a** | **CRE** |  |  |  | 9.2 ± 1.7 (2.6) |  |  |
|  | **SRE** |  |  |  | 1.6 ± 2 (4.1) |  |  |
|  | **Gqi** |  |  | 31 ± 1.5 (2.5) | 0.0009 ± 1.4 (3.2) |  |  |
| **SPX1b** | **CRE** |  |  |  | 6.4 ± 2.1 (2.8) |  |  |
|  | **SRE** |  |  |  | 5.2 ± 1.6 (4.1) |  |  |
|  | **Gqi** |  |  | 88 ± 1.3 (2.4) | 0.0003 ± 1.4 (3.2) |  |  |
| **Galanin** | **CRE** |  |  | 1.7 ± 2.5 (2.8) |  |  |  |
|  | **SRE** |  |  |  | 353 ± 2.1 (3.7) |  |  |
|  | **Gqi** |  |  | 0.0001 ± 5.6 (2.5) | 68 ± 1.3 (2.9) |  |  |

cAMP responsive element (CRE)-Luc was used as a reporter of PKA activation; Serum responsive element (SRE)-Luc was used as a reporter of PKC activation. Results are expressed as mean ± SEM and maximum fold induction value in brackets. Only values >2 presented.
