## Supplementary Figures 1-4 for "Spexin and a novel cichlid-specific spexin paralog both inhibit FSH and LH through a specific galanin receptor (Galr2b) in tilapia"

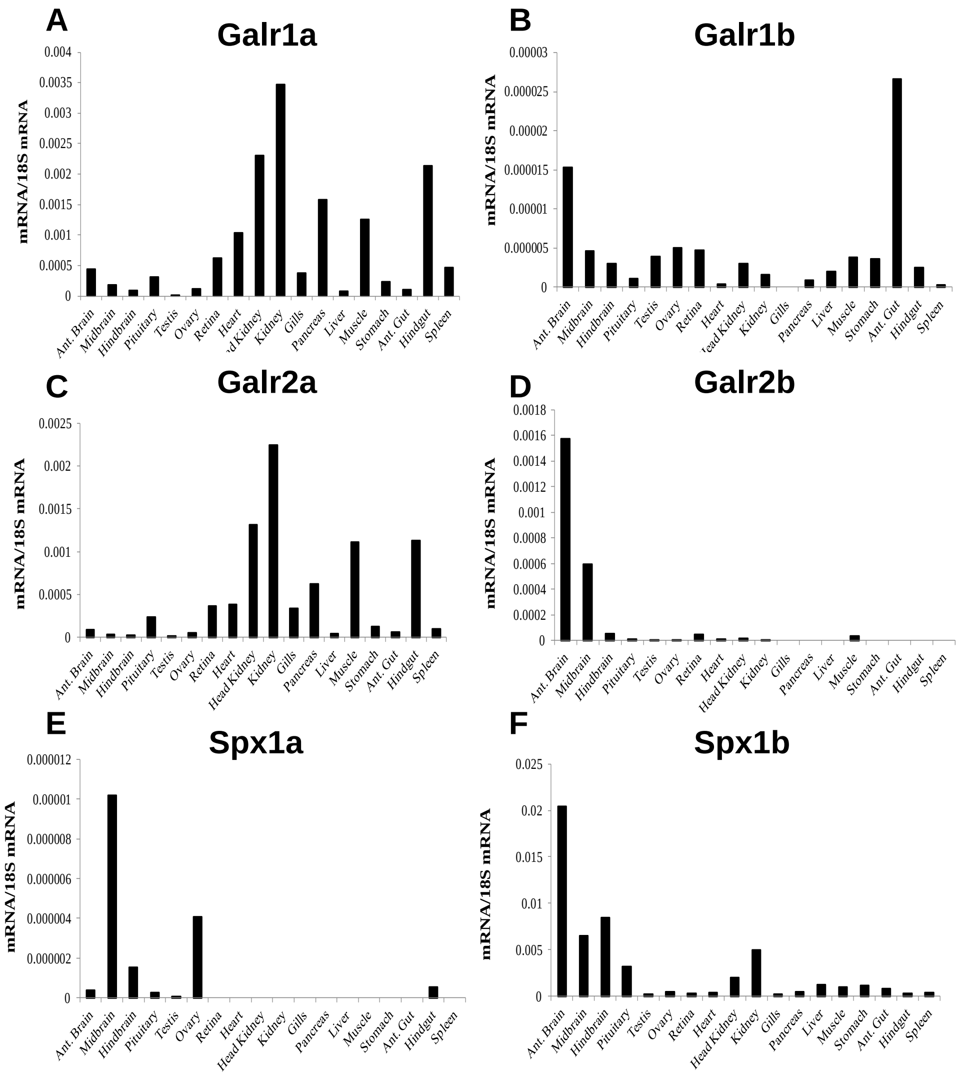


**Supplementary figure 1. Tissue distributions of tilapia SPXs and GALRs.** The genes were normalized against an endogenous reference (18S) by the comparative cycle threshold method.

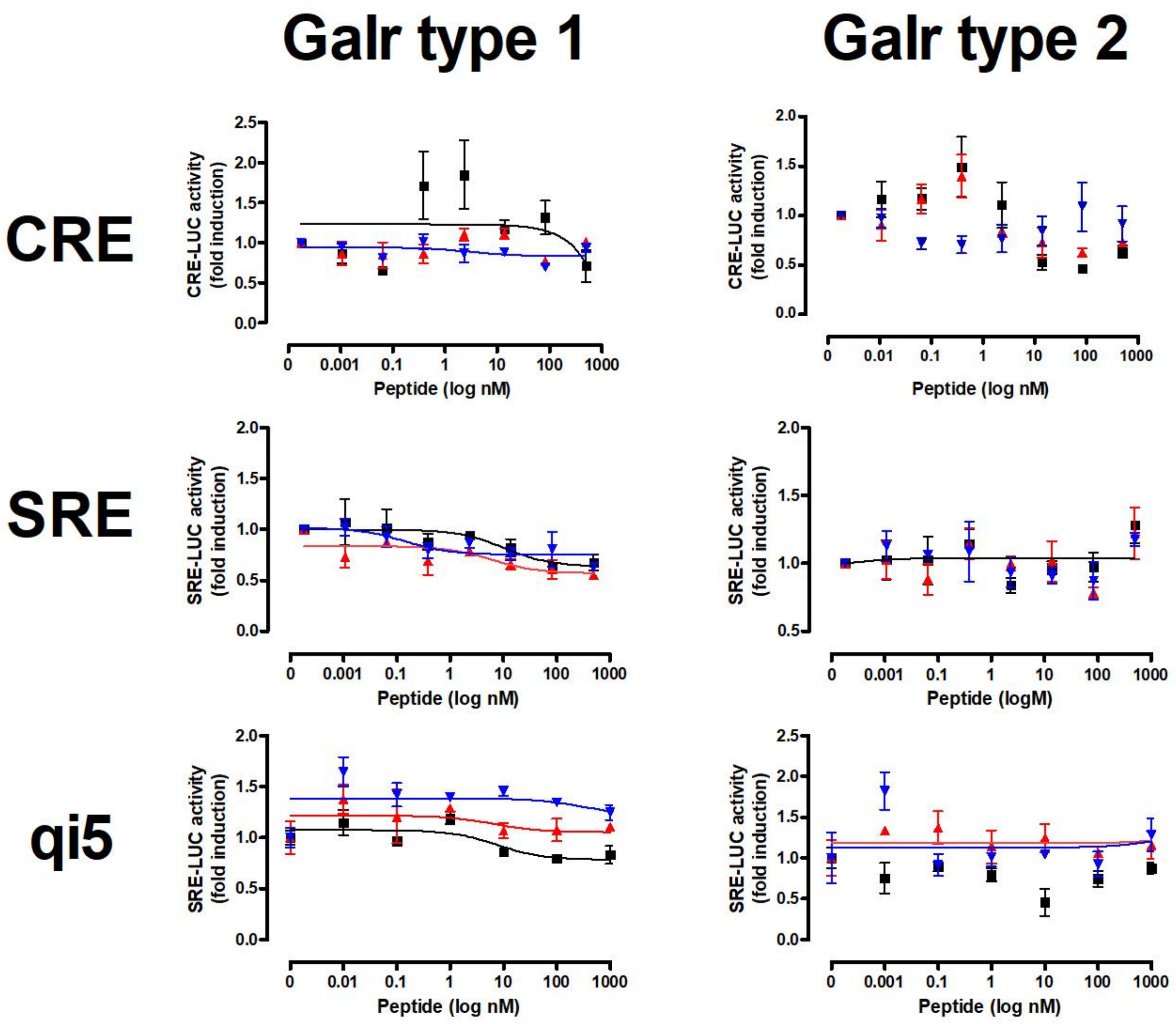


**Supplementary figure 2. In vitro reporter assays for tilapia Galanin receptors.** Tilapia Galr type 1 or type 2 were cotransfected with either CRE luc , SRE Luc, or Gqi5 into Cos7 cells and treated with either Spx1a (black squares), Spx1b (red triangles), or human galanin (Gal; blue upside-down triangles). Data are represented as mean ± SEM fold change in luciferase activity over basal (no treatment).

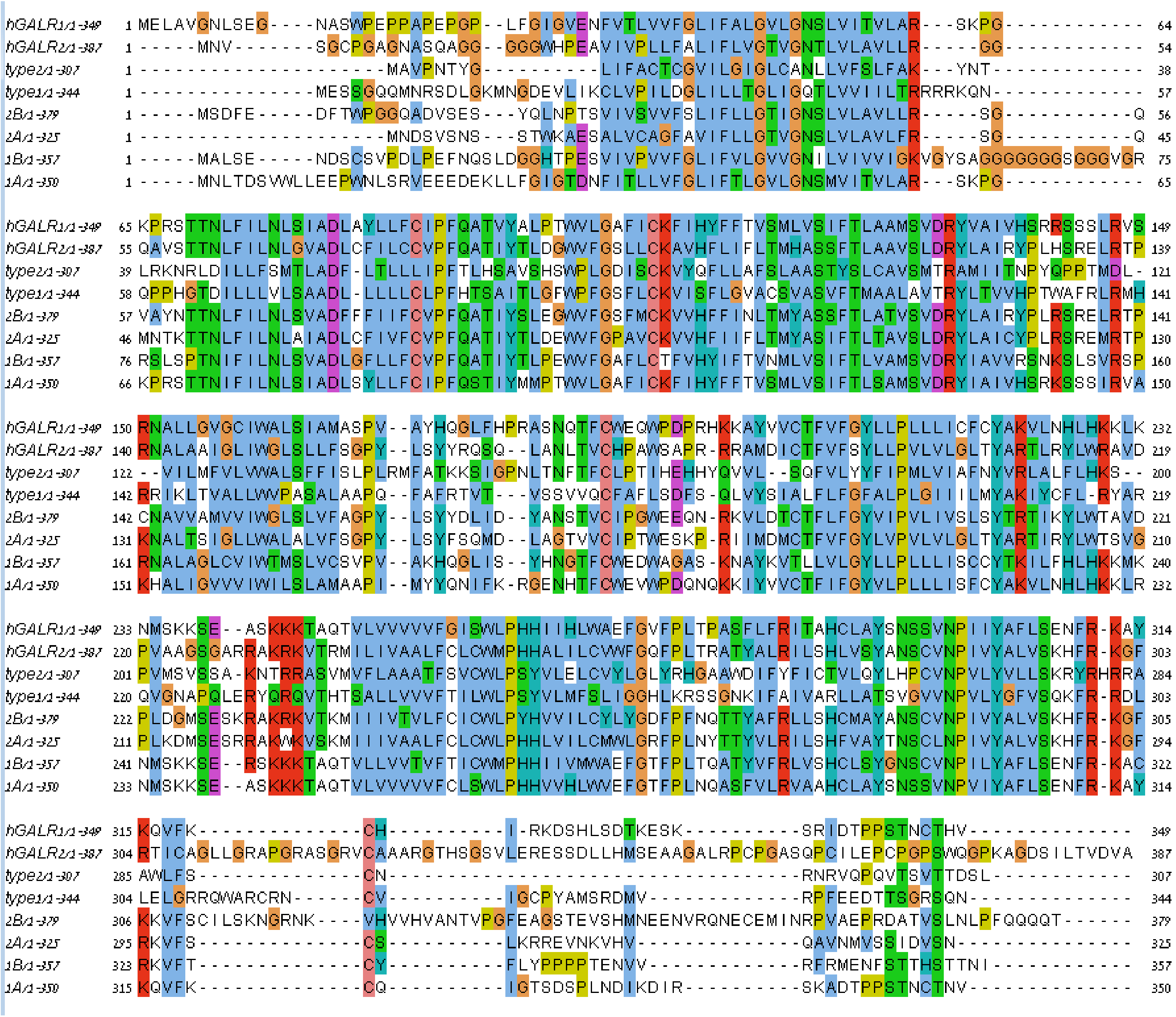


**Supplementary figure 3. Multiple sequence alignment of Galanin receptors**. hGALR1, hGALR2, Galr1a, Galr1b, Galr2a, Galr2b, Galr type 1, and Galr type2 amino acid sequences were aligned by MUSCLE, visualized in Jalview, and colored according to ClustalW conservation.

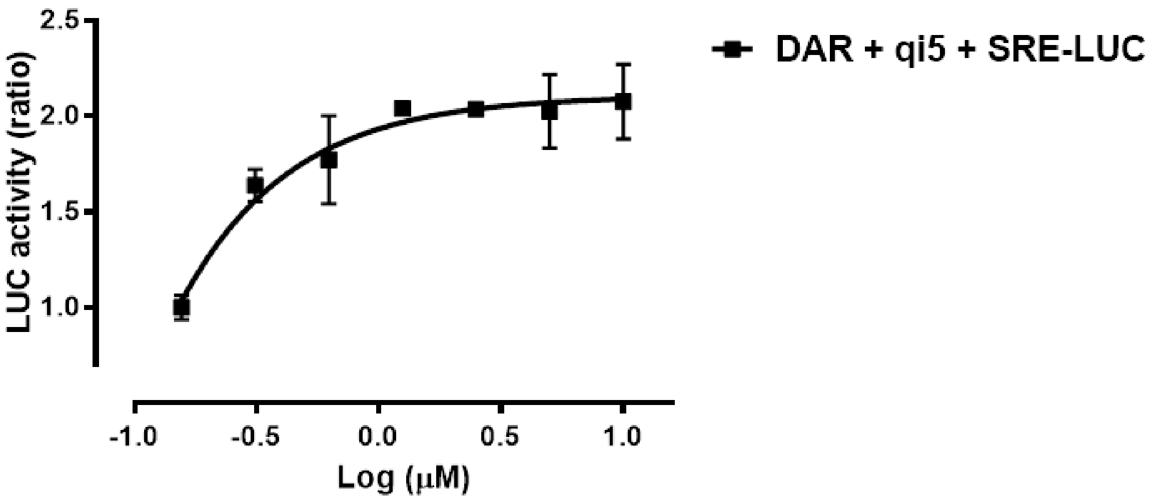


| | |
|---|---|
| EC50 | ~ 0.006253 |

**Supplementary figure 4.** ***In vitro*** **reporter assays for tilapia dopamine (D2) receptor.** Tilapia dopamine receptor was cotransfected with SRE-Luc and Gqi5 into Cos7 cells and treated with dopamine. Gqi5 allows inhibitory effects to be visualized as stimulus. Data are represented as mean ± SEM fold-change in luciferase activity over basal (no treatment).
